# Base-substitution mutation rate across the nuclear genome of *Alpheus* snapping shrimp and the timing of isolation by the Isthmus of Panama

**DOI:** 10.1101/2020.11.25.396556

**Authors:** Katherine Silliman, Jane L. Indorf, Nancy Knowlton, William E. Browne, Carla Hurt

## Abstract

The formation of the Isthmus of Panama and final closure of the Central American Seaway (CAS) provides an independent calibration point for examining the rate of DNA substitutions. This vicariant event has been widely used to estimate the substitution rate across mitochondrial genomes and to date evolutionary events in other taxonomic groups. Nuclear sequence data is increasingly being used to complement mitochondrial datasets for phylogenetic and evolutionary investigations; these studies would benefit from information regarding the rate and pattern of DNA substitutions derived from the nuclear genome. To estimate this genomewide neutral mutation rate (μ), genotype-by-sequencing (GBS) datasets were generated for three transisthmian species pairs in *Alpheus* snapping shrimp. Using a Bayesian coalescent approach (G-PhoCS) applied to 44,960 GBS loci, we estimated μ to be 2.64E-9 substitutions/site/year, when calibrated with the closure of the CAS at 3 Ma. This estimate is remarkably similar to experimentally derived mutation rates in model arthropod systems, strengthening the argument for a recent closure of the CAS. To our knowledge this is the first use of transisthmian species pairs to calibrate the rate of molecular evolution from GBS data.

## Introduction

The rate of DNA substitution is an essential parameter in evolutionary biology because it is used to establish a timeline for the history of life. In the field of phylogeography, molecular clocks have been applied extensively, as they enable investigators to put absolute values on measures of interest such as timing of speciation, patterns of historical migration, and estimates of effective population sizes (Cunningham and Collins 1994; Takahata et al. 1995; Arbogast et al. 2002; Bromham and Penny 2003; Obbard et al. 2012). Estimates of DNA substitution rates can be calibrated using experimental approaches (Baer et al. 2007), or by associating molecular phylogenies with independent information regarding the timing of species divergence (Ho and Duchêne 2014). The fossil record has been the most widely used source for calibrating rates of molecular evolution; however, in groups that lack a good fossil record, well-dated biogeographic barriers can be used for establishing the timing of species divergence (Ho et al. 2015).

The final closure of the Central American Seaway (CAS) and formation of the Isthmus of Panama provides a useful calibration point for examining the rates and patterns of molecular evolution, as the completion of the Isthmus of Panama created a nearly impenetrable barrier to gene flow for thousands of marine taxa. The splitting of multiple independent populations is particularly useful for molecular clock calibrations because it provides both absolute rates of divergence, and critical information regarding the constancy of molecular evolution rates across independent evolutionary lineages (Hickerson et al. 2010). By far the most cited transisthmianbased molecular clock calibrations come from the snapping shrimp genus *Alpheus* (Knowlton and Weigt 1998; Lessios 2008; Hurt et al. 2009). *Alpheus* contains more transisthmian species pairs than any other genus studied to date, providing a naturally replicated data set ideal for testing evolutionary hypotheses. However, comparisons of genetic distance estimates at mitochondrial genes across this genus have been shown to vary more than fourfold (Knowlton and Weigt 1998), which could be due to irregularity of the molecular clock or non-simultaneous divergence of transisthmian sister species. The latter explanation has been supported by multiple lines of evidence, including concordance of mitochondrial divergences with patterns of mating incompatibility and estimates of divergence from protein electrophoresis (Knowlton et al. 1993). Previously, Hurt et al. (2009) used a Bayesian coalescent approach to test simultaneous divergence of eight alpheid sister species using population-level sampling of multi-locus nuclear and mitochondrial genes. This work identified five transisthmian species pairs for which molecular data was consistent with recent and simultaneous divergence as a result of the closure of the CAS; these taxa are thus particularly well-suited for examining patterns of molecular divergence across the Isthmus of Panama.

The formation of the Isthmus is one of the most well-studied biogeographic vicariant events. Studies based on Foraminifera, isotope ratios, molecular phylogeography, and fossils suggest completion of the Isthmus had occurred by 3.5-2.7 Ma (Keigwin 1982; Schmidt et al. 2007; Lessios 2008; Molnar 2008; Coates and Stallard 2013; Jackson and O’Dea 2013; O’Dea et al. 2016), although there has been some recent debate about this conclusion. For example, some geological work has suggested an earlier seaway blockage, where the primary closure of the CAS occurred before 10 Ma, with only minor connections after that time via narrow, transient, shallow channels (Montes et al. 2012, 2015; Sepulchre et al. 2014; Jaramillo et al. 2017). Bacon et al. (2015) supported this earlier formation date using molecular and fossil data to determine that an initial land bridge was present 23-25 Ma and formation of the Isthmus occurred between 10 and 6 Ma. However, the assumptions and methods underlying these studies have been challenged, in particular, the inappropriate application of a universal rate of mitochondrial DNA divergence across clades and failure to account for ancestral lineage sorting (Lessios 2015; Marko et al. 2015; O’Dea et al. 2016) can provide more robust estimates of divergence times and inform understanding of the timing of the closure of the CAS (Bacon et al. 2015b).

The vast majority of transisthmian molecular clock calibrations have been applied to nucleotide sequence data from mitochondrial genes (Lessios 2008; Lavinia et al. 2016). However, improvements in DNA sequencing technology and increased awareness of the limitations of single locus mitochondrial data sets have transformed the fields of population genetics and phylogeography (Garrick et al. 2015). Reduced representation sequencing techniques, such as genotyping-by-sequencing (GBS), systematically target a subset of the genome by relying on restriction enzymes and shared cut sites. These techniques have proven to be useful and cost-efficient methods for screening polymorphisms at thousands of loci without the need for a reference genome (Elshire et al. 2011; Andrews et al. 2016). However, little is known about the rate and variance of nuclear DNA substitutions across GBS loci. Estimates of demographic parameters (e.g., effective population size, migration rates) from GBS data would benefit from a GBS-derived mutation rate (μ), as μ is often required to calculate absolute parameter estimates (Excoffier et al. 2013).

Inherent characteristics of GBS methodologies, including bioinformatic processing and the often large, nonrandom proportion of missing data, have the potential to bias demographic analyses (Eaton et al. 2017; O’Leary et al. 2018). Because these methods utilize restriction enzymes to reduce the genome, they require conservation of enzyme recognition cut sites to recover shared data among individuals. This typically results in a non-uniform distribution of reads across loci and individuals, and thus a reduced set of loci shared across all individuals (Eaton et al. 2017). The proportion of shared restriction sites (and sequenced loci) across sampled individuals is expected to decline as divergence times increase. Conserved regions of the genome, possibly regions under purifying selection, will be disproportionately represented when filtering criteria are strict, while faster evolving, neutral regions may be filtered out. This pattern has important implications for optimizing filtering parameters in bioinformatics pipelines. Many GBS/RAD papers have taken a conservative approach and employed strict filters for missing data (Campagna et al. 2015). However, *in silico* and empirical work have begun to show that a “total evidence approach” including loci with missing data is acceptable and may even be preferable in phylogenetic and population genetic studies (Huang and Knowles 2016; Eaton et al. 2017; Shafer et al. 2017; Tripp et al. 2017). Empirical investigations examining the influence of filtering criteria on estimates of demographic parameters would be useful for optimizing bioinformatic pipelines.

Here we report results from a comparative genomic study utilizing *Alpheus* species pairs to examine patterns of molecular divergence across the nuclear genome. Reduced representation GBS datasets were generated for three transisthmian species pairs: *A. malleator/A. wonkimi, A. formosus/A. panamensis,* and *A. colombiensis /A. estuariensis.* First, we investigated the phylogenetic signal of shared GBS-derived sequence tags in order to identify potential sequence bias due to divergence of restriction sites. The optimized GBS dataset was then used to estimate the timing of divergence (τ) for the selected species pairs using a Bayesian coalescent modelling approach while correcting for variance in divergence times. We then estimated the rate of base-substitutions (μ) using the final closure of the CAS as a calibration point. In order to evaluate claims of an earlier closure of the CAS, both 3 Ma and 10 Ma were used as calibration points for calculating μ, and the results were then compared to estimates of substitution rates in other multicellular eukaryotes. To our knowledge, this is the first use of transisthmian species pairs to calibrate the rate of molecular evolution across the nuclear genome.

## Methods

### Sample Collections

Three transisthmian *Alpheus* sister species pairs (six species total) were selected for GBS sequencing and analysis: the eastern Pacific/western Atlantic pairs *A. malleator/A. wonkimi, A. colombiensis/A. estuariensis,* and *A. panamensis/ A. formosus* (Table 1). Previous work suggested that divergence times for these taxa were contemporaneous and likely to have resulted from the final closure of the Isthmus (Hurt et al. 2009). All shrimp were collected from the Caribbean and Pacific coasts of Panama. *Alpheus panamensis* and *A. formosus* were collected from intertidal or subtidal habitats, *A. malleator* and *A. wonkimi* from exposed shores, burrowed inside crevices in hard substrate, and *A. colombiensis* and *A. estuariensis* were collected from mudflats near mangroves. All samples were frozen in liquid nitrogen and stored at −80 °C. DNA sequences from the mitochondrial gene cytochrome oxidase I (COI) were generated for all included individuals and compared to previously recorded COI sequences from the corresponding species. Primers and PCR conditions for amplification of COI followed Hurt et al. (2013).

**Table 1.** Collection information for Alpheus samples used for GBS sequencing including sample size (N), distribution (EP = Eastern Pacific, WA = Western Atlantic), known habitat, and Genbank Accession numbers. ^1^Alpheus wonkimi was referred to as A. cf malleator or A. isthmalleator in Hurt et al. (2009).

| Species | N | Distribution | Habitat | COI Genbank Accessions |
| --- | --- | --- | --- | --- |
| *A. panamensis* | 5 | EP | Intertidal under rocks and coral rubble |  |
| *A. formosus* | 5 | WA | Intertidal under rocks and coral rubble |  |
| *A. colombiensis* | 4 | EP | Burrows in mangroves |  |
| *A. estuariensis* | 4 | WA | Burrows in mangroves |  |
| *A. wonkimi*<sup>1</sup> | 4 | EP | Endolithic in rock crevices |  |
| *A. malleator* | 2 | WA | Endolithic in rock crevices |  |

### Molecular Methods

Genomic DNA was extracted from 24 individuals using the DNeasy tissue kit (Qiagen Inc., Valencia, California), and samples were treated with RNase following the manufacturer’s protocol. Three to four replicate GBS libraries per individual were optimized, generated, and sequenced at the Cornell University Biotechnology Resource Center Genomic Diversity Facility following the protocol of (Elshire et al. 2011), resulting in a total of 96 samples. Briefly, genomic DNA was digested with EcoT22I (A|TGCAT) and barcoded adapters were ligated onto resulting restriction fragments. Pooled libraries were sequenced on a single Illumina HiSeq 2000/2500 lane, obtaining 100 base pair, single-end sequencing reads. Sequence reads from replicate libraries were combined for downstream analyses.

### Quality Filtering, Locus Assembly, and Genotyping

Raw sequencing reads were demultiplexed, quality filtered, and *de novo* clustered using pyRAD v.3.0.2 (Eaton 2014), a pipeline optimized to produce aligned orthologous loci across distantly related taxa using restriction-site associated DNA. Demultiplexing used sample-specific barcode sequences, allowing one mismatch in the barcode sequence. Base calls with a Phred quality score under 20 were converted to Ns, and reads containing more than 4 Ns were discarded. Adapter sequences, barcodes, and the cut site sequences were trimmed from reads passing filter, with only reads greater than 50 bp retained. For within-sample clustering, a minimum coverage cutoff of 5× was employed. Consensus sequences with more than eight heterozygous sites were discarded as potential paralogs. Clustered orthologs containing heterozygous sites that were shared by more than two samples were also discarded as putative paralogs. The same clustering threshold of 85% was used for both within- and across-sample clustering (Eaton 2014).

We generated 11 datasets that varied in included samples (10–24), the minimum number of samples (*m*) that had to be shared by each locus (3–6), and the minimum number of species (*s*) shared by each locus (1–3). In particular, one *A. malleator* individual had very few sequencing reads and therefore was excluded from some datasets. These additional datasets were used to investigate the impact of filtering by sample coverage and missing data on estimates of demographic parameters (Supp. Table 1). The primary dataset, Am4s2, used in the following analyses includes all samples (A), with at least four individuals (m4) and two species (s2) genotyped at each locus. Transition/transversion (Ts/Tv) ratios were calculated for all datasets using VCFtools (Danecek et al. 2011).

### Phylogenetic Analyses

For phylogenetic reconstructions, a concatenated matrix was produced for the Am4s2 dataset and partitioned into neutral and coding sites, with model parameters estimated independently for each partition. Maximum Likelihood inferences were conducted using RAxML v8.1 (Stamatakis 2014) under the General Time-Reversible nucleotide model with gamma-distributed rate heterogeneity (GTRGAMMA) and 1000 bootstrap replicates. Bayesian inferences were performed using Exabayes v1.4 (Exelixis Lab, http://sco.hits.org/exelixis/web/software/exabayes/). Four independent MCMC runs were run for 1,000,000 generations, sampling every 500 generations. Runs were initiated from a random order addition parsimony tree. All other settings were default. Pairwise branch length distance between each species was calculated using the *ape* package in R (Paradis et al. 2004).

### Locus Bias

To assess patterns of locus-sharing among individuals, the R package RADami (Hipp et al. 2014) was used to construct a locus presence-absence (LPA) matrix and display the proportion of shared loci between pairs of individuals (Fig. 3a). Pairwise Jaccard’s distances calculated from this matrix were visualized using nonmetric multidimensional scaling in the R package vegan (Oksanen et al. 2016). The dimensionality of the ordination was determined by performing 50 replicate runs at random starting configurations for K = 1 to 10 axes, with K = 1 to 3 showing the largest decreases in final stress. Both the K = 2 and K = 3 ordinations were rerun with 2000 replicates each, and converged on best solutions. Only the K = 3 ordinations are reported in this study, as they provided the clearest visualization of sample clustering. The poorly sequenced *A. malleator* individual was excluded from ordinations because low overlap in locus coverage between this individual and all others dominated the ordinations in preliminary analyses (not shown).

To understand the influence of phylogenetic distance and sequencing depth on locus bias, we built linear models of pairwise shared loci across all samples using the *lm* function in base R. First, we constructed a 24 × 24 matrix of the number of pairwise shared loci between all samples using the pyRAD .loci file, based on Python and R code from (Winston et al. 2016). We then used a linear model to predict the number of shared loci between two samples based only on the combined number of reads after quality filtering. This model was compared to a linear model where filtered reads and phylogenetic distance between samples were the independent variables. Phylogenetic distances were determined from the RAxML maximum likelihood tree using the R package *ape* v3.5 (Popescu et al. 2012).

### Estimating Homology to Coding Regions

In order to identify putative protein-coding loci that may be subject to selection, a comprehensive dataset including all individuals and loci genotyped in at least 4 individuals (Am4) was blasted against Metazoa sequences in both the NCBI remote BLAST nucleotide database (nt) and protein database (nr), using the programs blastx and blastn in BLAST+ 2.3.0 (ftp://ftp.ncbi.nih.gov/blast/executables/LATEST/). Analyses used default settings and an “*E*-value” significance threshold of 1. These results were imported into Blast2Go, where InterProScan was used to identify putative protein coding sequences. Loci with significant matches to mRNA sequences, proteins, or matches to an InterPro database were classified conservatively as putative nonneutral loci. A local BLAST database was created with these loci and used to identify nonneutral loci in the other datasets, including Am4s2. Loci with InterPro matches or BLAST results with an “*E*-value” less than 1E-3 were annotated with Gene Ontology (GO) terms in Blast2Go (Götz et al. 2008) (Supp. Fig. 1).

### Demography, Gene Flow, and Mutation Rates

Estimates of demographic parameters, postdivergence gene flow, and genome-wide mutation rate were performed using the Generalized Phylogenetic Coalescent Sampler (G-PhoCS) version 1.2.3 (Gronau et al. 2011), which infers divergence times (τ), ancestral effective population sizes (Θ), and migration rates. A Markov Chain Monte Carlo (MCMC) sampling strategy was used to sample parameters from a full coalescent isolation-with-migration model, where post-divergence migration bands are optional and specified by the user. This model assumes a separate constant population size for each branch of the phylogeny, and a separate constant migration rate for any migration bands specified. All demographic parameters are scaled by the mutation rate (μ), which can either be held constant or allowed to vary across loci. G-PhoCS takes as input a given phylogeny, specified directional migration bands, and a collection of aligned neutrally evolving loci, where heterozygous genotypes are unphased and the likelihood computation analytically sums over all possible phasings.

Due to the computationally intensive nature of this analysis, we were unable to analyze the full Am4s2 neutral dataset (44,960 total loci). Thus the full dataset was reduced to 14,986 randomly sampled loci. Analyses were initially performed under the assumption of no gene flow after divergence, estimating 16 parameters (11 population sizes and 5 divergence times). Three replicate analyses were conducted with mutation rate held constant, as well as an additional three analyses with random locus-specific mutation rates estimated by G-PhoCS. As mutation rates are known to vary across genomes, we expected the latter analyses to provide a more accurate estimation of overall mutation rates. All MCMC runs were executed using the same settings, unless otherwise indicated. Each Markov chain included 100,000 burn-in iterations, after which parameter values were sampled every 10 iterations for 200,000 iterations. The prior distributions over model parameters were defined by a product of Gamma distributions (Supp. Table 2). The fine-tune parameters of the MCMC procedure were set automatically during the first 10,000 burn-in iterations (using the ‘find-finetunes TRUE’ option in the G-Phocs control file). We conditioned on the phylogenetic relationships of taxa based on the ML tree and phylogenetic inference in (Williams et al. 2001). Convergence for each run was inspected manually in Tracer (Rambaut et al. 2018). Replicate runs were combined for calculating the mean and confidence intervals of parameter estimates.

We conducted multiple GPHoCS runs on ten datasets that varied in taxa composition and the minimum number of individuals/species recovered at a locus to explore the influence of filtering by sample coverage on parameter estimates. The purpose of this approach was to 1) determine how different stringency filters for missing data affected demographic estimates, and 2) ensure that demographic estimates were robust to the selection of species included in the analysis. Three of the datasets included representatives from all six species, and one dataset only included two species pairs (PFECm3s2). The other six datasets contained three species each— one transisthmian species pair and one outgroup species. These triplet datasets were classified into an *a* group and a *b* group for visualization in Figure 4. In total, τ and Θ were estimated across 23 runs for *A. estuariensis/A. colombiensis* (τ_EC_, Θ_EC_) and *A. panamensis/A. formosus* (τ_PF_, Θ_PF_) and 21 runs for A. wonkimi A. malleator (τ_WM_, Θ_WM_) (Supp. Table 1).

We also conducted G-PhoCS analyses that allowed for migration between sister species in order to explicitly test for post-divergence gene flow and determine the effect of gene flow on estimates of population divergence times and effective population sizes. Three replicate runs on the Am4s2 dataset included 6 directional ‘migration bands’ representing gene flow between each sister species pair, with the mutation rate (μ) allowed to vary across loci. Following (Freedman et al. 2014), a migration band was inferred to have significant gene flow if the 95% Bayesian credible interval of the migration rate *(M)* did not include 1E-5 in any of the replicate runs. We then conducted three replicate G-PhoCS analyses incorporating the migration bands between the sister species pair that showed significant gene flow. The effective number of migrants per generation was calculated as *M*_A→B_ × Θ_B_.

We used the outputs of the three replicate G-PhoCS analyses with random locus-specific mutation rates that incorporated the migration bands between the one sister species pair that showed significant gene flow to obtain a best estimate of τ and Θ for each species pair. MCMC samples were combined from the posterior distributions for the replicate runs to determine the mean and 95% confidence interval estimates for the demographic parameters. The timing of species divergences (τ) estimated by G-PhoCS were calibrated with estimates of the final closure of the CAS to estimate the absolute rate of μ and its variation among sister species pairs. Previous work has shown that these three species pairs have contemporaneous divergence times (within 5 my); therefore, we divided the τ estimated by G-PhoCS for each species pair by a similar absolute divergence time to obtain estimates of μ. We estimated μ using both 3 Ma and 10 Ma as calibration points, as the final closure of the Isthmus has been proposed to occur within this time interval.

## Results

### Sequence Assembly and Gene Ontology

Sequencing of 24 individuals across 96 libraries yielded 189,408,593 total raw sequencing reads (average of 7,350,671 ± 3,318,208 reads per sample). After quality filtering, replicate samples were combined to assemble consensus sequences for each individual, with a mean read depth of 14.33 (± 63.42). These were further filtered to 43,831 ± 25,944 consensus sequences per individual. Datasets differing by sample coverage or included taxa varied in the total number of loci (3,481-48,062), Ts/Tv ratios (1.359-1.594), and the amount of missing data (Supp. Table 1). In neutral loci from the Am4s2 dataset, the frequency of C/G substitutions was less relative to other substitutions (Fig. 1).

**Figure 1.**
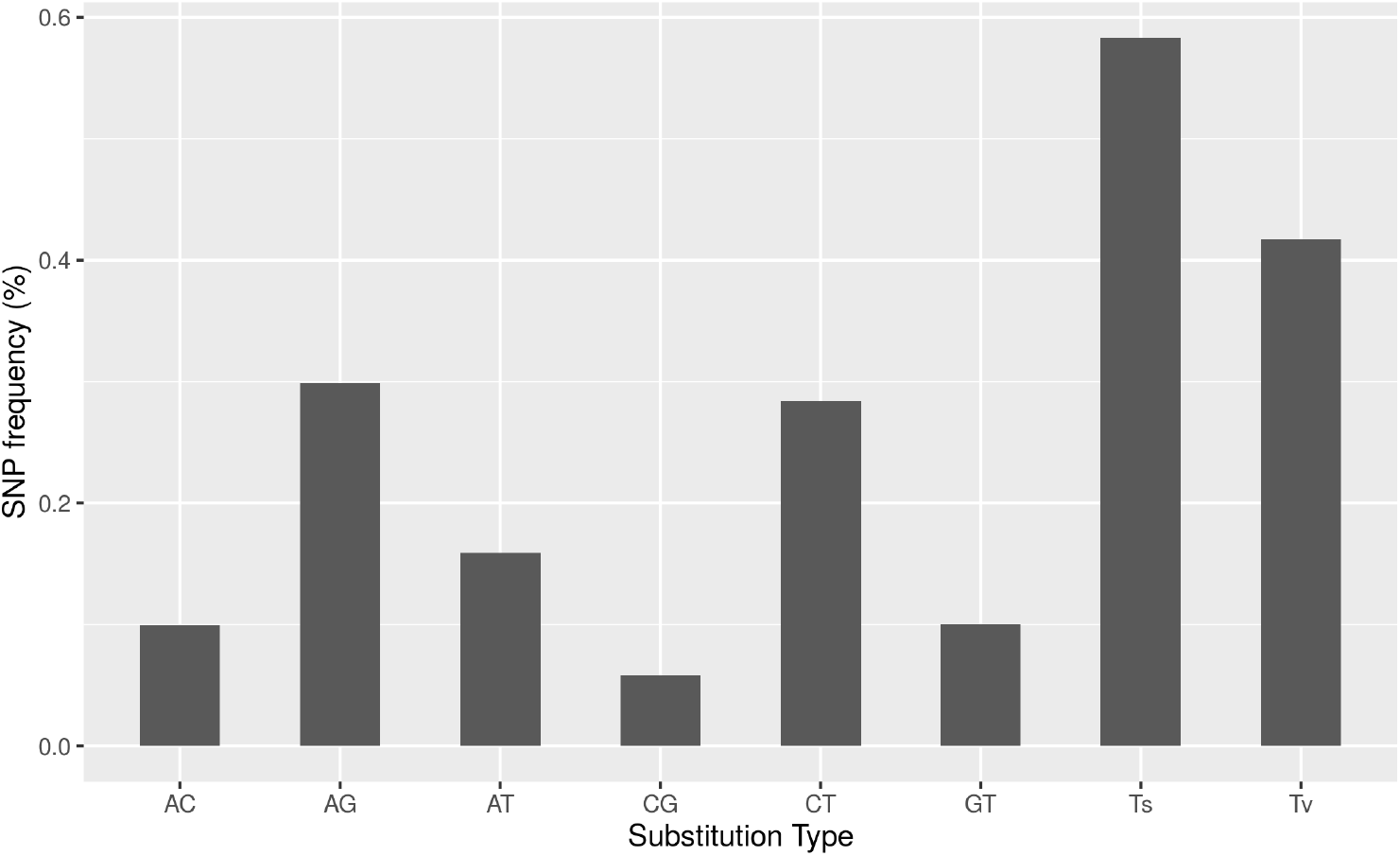
Distribution of six types of substitutions based on 222,174 SNPs across 44,960 loci in the Am4s2 dataset, after filtering for putative neutral loci.

Of the 56,838 loci in the Am4 dataset, 3,925 loci (6.9%) were identified as non-neutral based on inferred homology to mRNA or protein coding sequences using BLAST tools and InterProScan. Whiteleg shrimp *(Litopenaeus vannamei)* had the most top hits in the nr database; 707 loci had a hit to InterProScan and 647 loci had a Blast E value of 0.001 to the nr or nt databases, of which 261 were annotated with GO terms (Supp. Fig. 1).

### Locus Bias and Phylogenetics

Phylogenetic reconstructions resulted in monophyletic species and sister-species pairs with 100% bootstrap support, and *A. estuariensis* and *A. colombiensis* clustered separately from the other four species (Fig. 2). These topologies are consistent with previous phylogenetic work based on the mitochondrial gene cytochrome oxidase I (COI) and two nuclear genes (Williams et al. 2001). Ordination of the pairwise shared-locus matrix showed a strong phylogenetic signal, with individuals from the same sister species pair clustering together (Fig. 3). This result is consistent across datasets varying in sample coverage (Supp. Fig. 2). Of the two linear models tested, our model that included phylogenetic distance and the log-root product of total number of reads passing filter performed the best [r^2^ = 0.872, p=0] (Supp. Fig. 3).

**Figure 2.**
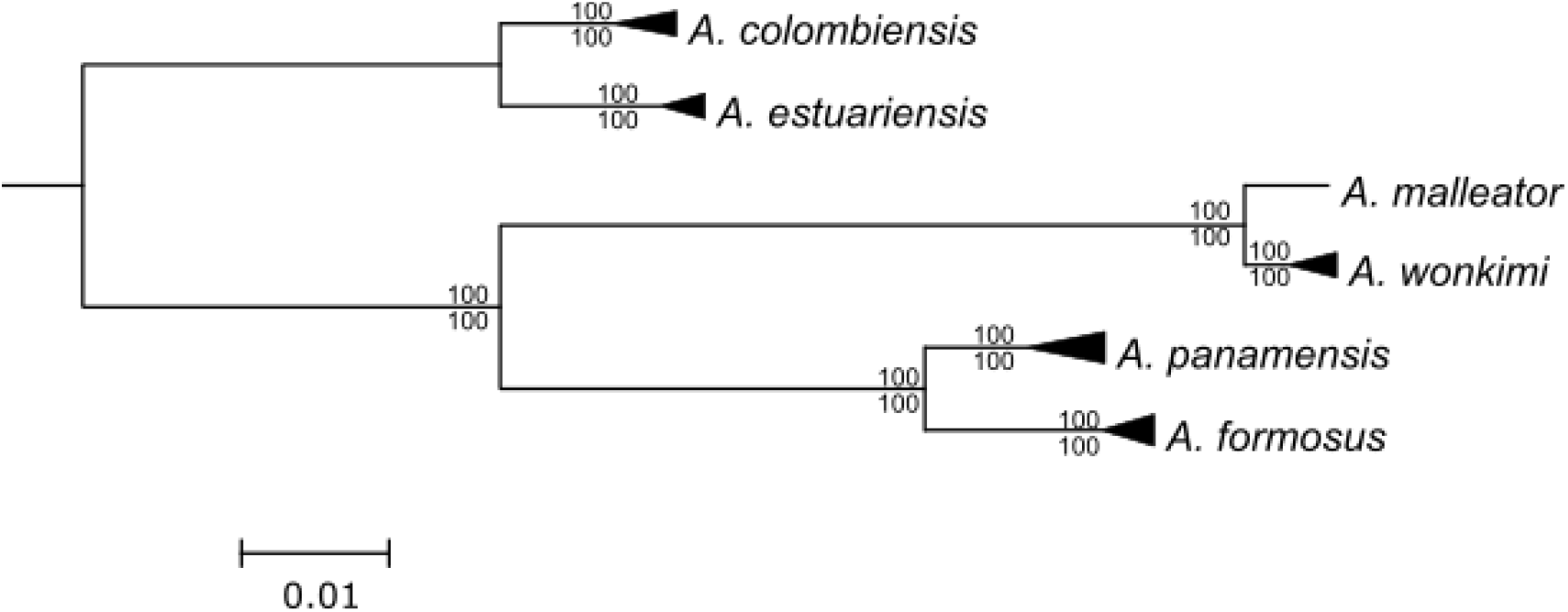
Results from Bayesian phylogenetic reconstructions based on concatenated matrices from the full dataset (23m6s). Nodes were collapsed by species. All species and transisthmian sister-species pairs were monophyletic with 100% posterior probability support (values above node). Maximum likelihood reconstructions using the same dataset resulted in an identical topology with 100% bootstrap support for every node (values below nodes).

**Figure 3.**
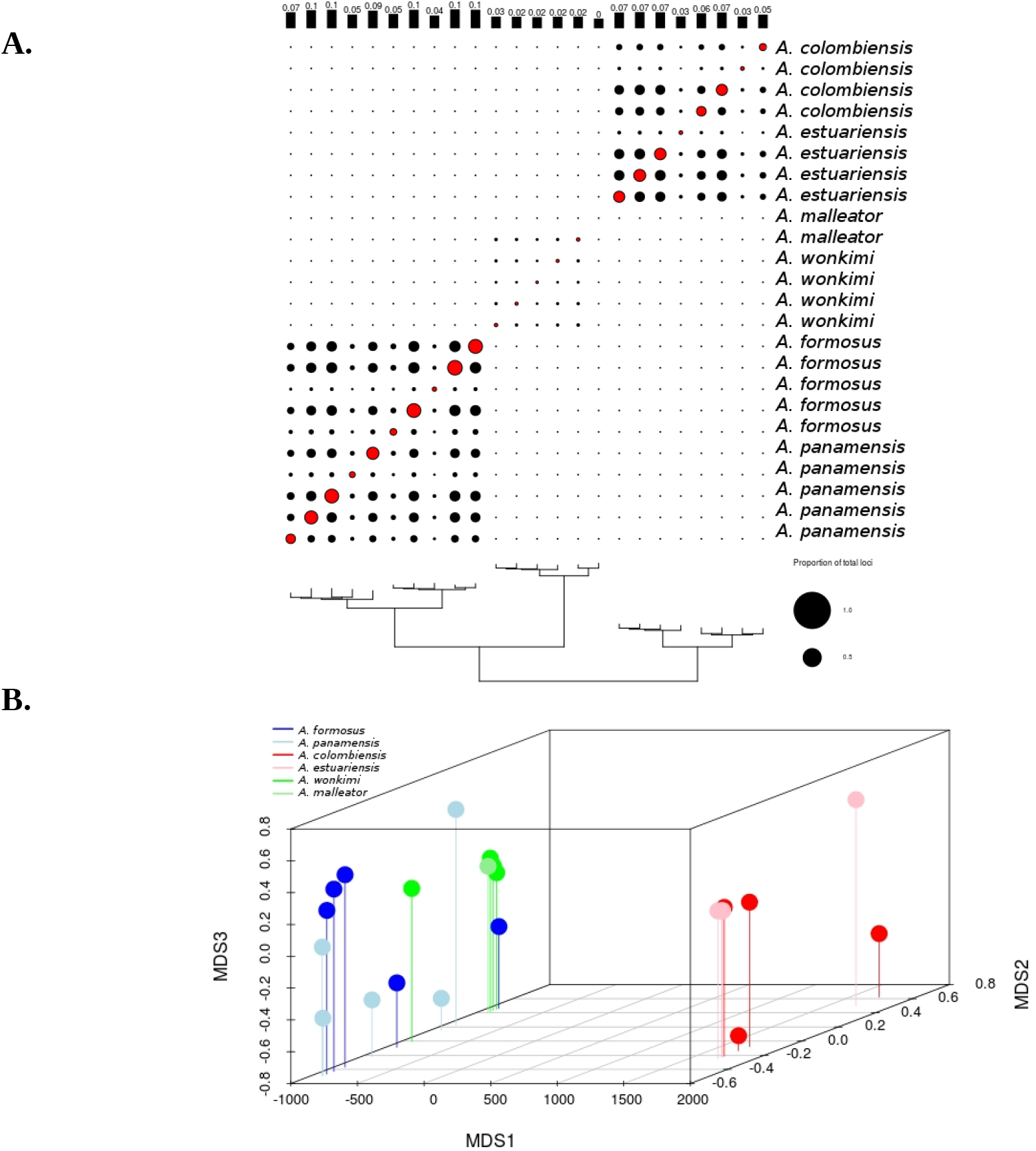
a) Proportion of loci shared among individuals in the Am4s2 dataset. Loci shared between individuals (black circles) or successfully amplified within an individual (red circles) expressed as the proportion from 0 to 1 of all 48,062 loci scored in the Am4s2 dataset. Plotted with the phylogenetic tree based on maximum likelihood inference, with branch lengths scaled in substitutions per nucleotide. b) Ordination of *Alpheus* samples based on nonmetric multidimensional scaling of the locus presence-absence matrix, K=3, colored by species.

### Demographic inference and post-divergence gene flow

We conducted multiple G-PhoCS runs on datasets that varied in species composition and the minimum number of individuals/species recovered at a locus, as the number of loci shared between samples was shown to be influenced by phylogenetic distance. Replicate runs on the same dataset were nearly identical, indicating there were a sufficient number of simulations to achieve convergence. We found the greatest consistency in parameter estimation across runs on the Am4s2, 23m6s2, and species triplet datasets (Figure 4). Dataset Am3s3, which required at least three species to be sequenced at every locus and thus had the least missing data, produced significantly lower estimates for τ_root_, τ_PF_, and τ_EC_. The results from Am3s3 were similar to those from 23m6s2 for τ_WM_, but both were significantly lower than the results from Am4s2 and the species triplet datasets (Figure 4).

**Figure 4.**
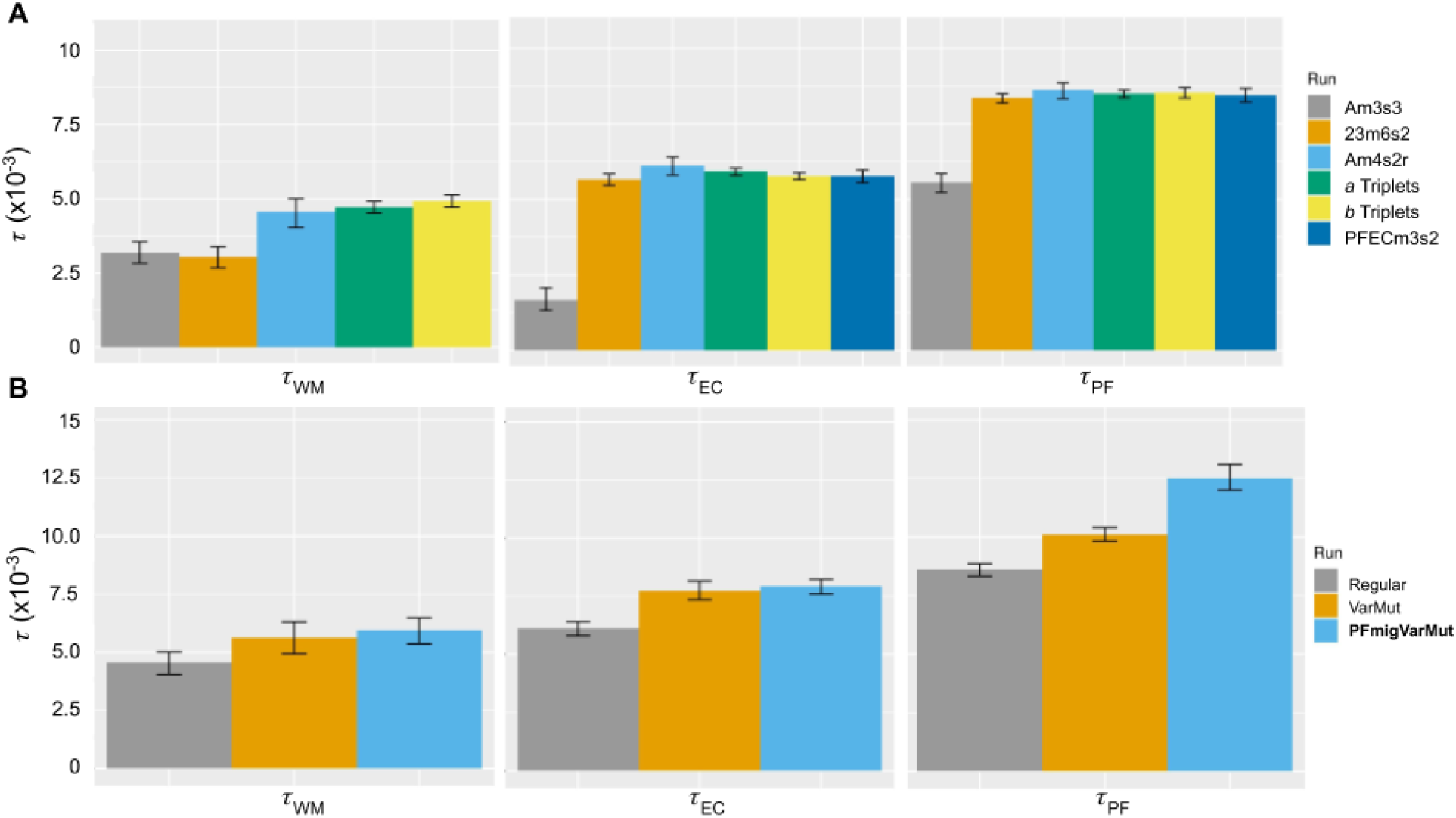
Estimates of divergence times (τ) for three transisthmian *Alpheus* species pairs (*A. wonkimi/A. malleator* (WM), *A. estuariensis/ A. colombiensis* (EC), *A. panamensis/ A. formosus* (PF)) across datasets, scaled by the substitution rate (μ). Error bars indicate 95% Bayesian credible intervals across all replicate runs. **A)** Datasets that varied in species composition and the minimum number of individuals (*m*) or species *(s)* recovered at a locus, modeled with a constant mutation rate across loci and no migration. Parameter values for each dataset are listed in Supp. Table 1. *a* and *b* triplet datasets include the focal species pair plus one outgroup species. **B)** The Am4s2r dataset, modeled with either constant μ across loci and no migration (Regular), variable μ across loci (VarMut), or variable μ and migration allowed between *A. panemensis/A. formosus* (PFmigVarMut).

The Am4s2 dataset was chosen for downstream analysis as it represented all taxa and gave consistent results. We conducted three G-PhoCS run replicates on Am4s2, allowing the rate of mutation to vary randomly across loci. This represents a more realistic scenario than simply estimating a single mutation rate for all loci; however, it increases the computational time (by at least 10%). In the original G-PhoCS paper, the authors determine through simulations and empirical analyses that a “random rates” model can possibly influence the estimation of root population divergence and ancestral effective population size, but it likely has only a minor effect on divergence times of more recent branches in the phylogeny (Gronau et al. 2011). We found that allowing for mutation rate to vary across loci also widened the confidence intervals around our estimates of τ_root_, as well as increasing estimates of τ at all population splits and decreasing estimates of Θ. (Table 4, Supp. File 2).

Tests for post-divergence gene flow between transisthmian sister species suggested that, of the six migration bands tested, gene flow was only significant for the *A. panamensis* / *A. formosus* species pair. Migration rate estimates from G-PhoCS were significant across all replicate runs for this species pair. When G-PhoCS was run with variable mutation rates across loci and migration only allowed between *A. panamensis* and *A. formosus*, we found an expected number of 0.130 migrants per generation from *A. panamensis −> A. formosus* and an expected number of 0.044 migrants per generation from *A. formosus −> A. panamensis.* Tests for postdivergence migration were not consistently significant for the other two species pairs.

To obtain our best estimate of τ in order to calculate μ, we performed three replicate runs on the Am4s2 dataset allowing for migration only between *A. formosus* and *A. panamensis* and variable rates of mutation across loci. *A. malleator/A. wonkimi* had the smallest estimated divergence time (τ_WM_ ~5.95E-3) and therefore likely diverged most recently, followed by *A. estuariensis /A. colombiensis* (τ_EC_~7.93E-3), and *A. panamensis/A. formosus* (τ_PF_~12.5E-3) (Table 2). As *A. malleator* only had sequence data for one individual at the majority of loci, we calculated *μ* from τ_EC_. If we assume annual generation times and divergence at the proposed final closing of the Isthmus (3 Ma) we get a substitution rate of 2.64E-9 (2.53E-9 - 2.75E-9). If we use the more controversial closure calibration of 10 Ma, as is determined in (Sepulchre et al. 2014),supported by (Montes et al. 2015), and cited in (Bacon et al. 2015a), we get a substitution rate of 7.93E-10 (7.6E-10 - 8.24E-10).

**Table 2.** Demographic inferences based on G-PhoCS including effective ancestral population size (Θ), absolute effective ancestral population size in number of individuals (N_e_), divergence time (τ), absolute divergence time in millions of years (T), and calculated mutation rate (μ). Numbers in parentheses indicate 95% confidence intervals.

| Species Pair | $\Theta$ | $N_e$ ($\times 10^4$) | $\tau$ (E-3) | T ($\mu = 2.64$) | $\mu$ using 3 Ma (E-9) | $\mu$ using 10 Ma (E-9) |
| --- | --- | --- | --- | --- | --- | --- |
| *A. wonkimi*/<br>*A. malleator* | 0.018<br>(0.016,0.019) | 167<br>(153,182) | 5.95<br>(5.36,6.48) | 2.25<br>(2.03,2.45) | 1.98<br>(1.79,2.16) | 0.59 (0.54,0.65) |
| *A. colombiensis*/<br>*A. estuariensis* | 0.027<br>(0.025,0.028) | 152<br>(240, 264) | 7.93<br>(7.60,8.24) | 3<br>(2.88,3.12) | 2.64<br>(2.53,2.75) | 0.79 (0.76,0.82) |
| *A. panamensis*/<br>*A. formosus* | 0.0205<br>(0.019,0.022) | 194<br>(184, 205) | 12.5<br>(12.0,13.1) | 4.73<br>(4.55,4.96) | 4.17<br>(4.00, 4.37) | 1.25<br>(1.20,1.31) |

## Discussion

The field of evolutionary biology is rapidly transitioning from its reliance on a handful of mitochondrial loci to the incorporation of genome-wide sequence data for reconstructing evolutionary histories. Reduced representation methods for genome sampling, such as GBS, have seen widespread applications for genomic investigations involving non-model organisms (Wells and Dale 2018; Titus et al. 2019). Interpretation of these genomic datasets will require an understanding of the rate of DNA substitution across the nuclear genome. Molecular clock calibrations utilizing the well-examined closure of the Isthmus of Panama are among the most widely used parameters for dating cladogenic events (Cunningham and Collins 1994; Knowlton and Weigt 1998; Lessios 2008,Bacon et al. 2015a). The genus *Alpheus* includes more transisthmian sister species pairs than any other taxonomic group, facilitating the development of replicated datasets needed for robustly estimating mutation rate. Our GBS dataset included 2,844,991 bp from 44,960 neutral loci and represented three alpheid transisthmian species-pairs known to have diverged comparatively recently (Hurt et al. 2009).

Collectively, results from our study can inform other studies utilizing reduced representation sequencing for evolutionary investigations. Below we outline the implications of our results for 1) using such datasets to estimate mutation rates and reducing the role of locus bias in these analyses; and 2) understanding the evolutionary processes associated with the rise of the Isthmus of Panama, both the timing of divergence events and implications for the debate on when final closure occurred.

### Mutation Rate Estimates

Accounting for coalescence within ancestral populations and post-split migration, our best estimate for the per site mutation rate (μ), was 2.64E-9 (2.53E-9-2.75E-9) substitutions/site/year using the more widely accepted estimated time of 3 Ma for closure of the Isthmus. This estimate is largely consistent with mutation rate estimates obtained experimentally in other model arthropods (Keightley et al. 2014; Keith et al. 2016). A previous analysis of the nuclear mutation rate in transisthmian *Alpheus* pairs (Hurt et al. 2009) yielded estimates an order of magnitude lower than most experimentally derived rates and those reported here. This earlier study used Sanger sequencing data from eight nuclear genes (4457 bp) and employed a coalescent-based method (MCMCcoal) to estimate an average per site mutation rate across loci of 2.3E-10 substitutions/year. As this estimate was based on sequence data exclusively from protein coding regions, the reduced substitution rate was likely the result of purifying selection. By excluding putative protein coding genes from our data set, our new estimate, based on 2,844,991 bp, is more similar to experimental mutation rate estimates in other arthropods (e.g. 3.46E-9 in *Drosophila (Keightley et al. 2009),* 3.8E-9 in *Daphnia* (Keith et al. 2016). Both MCMCcoal and the method used in our study, G-PhoCS, estimate ancestral population sizes and population divergence times by comparing and integrating across genealogies at multiple neutrally evolving loci. Importantly, G-PhoCS extends the MCMCcoal model by allowing for gene flow between diverged populations and facilitating the use of unphased genotype data. The latter is often necessary for GBS data when phasing is not possible.

The popularity of mitochondrial markers for phylogenetic and evolutionary studies is largely due to their higher mutation rate compared to nuclear markers. Our understanding of the ratio of substitution rates in the mitochondrial genome relative to the nuclear genome (μ_mit_/μ_nuc_) has largely been based on observations in vertebrate taxa (Brown et al. 1979). Allio et al. (2017) analyzed μ_mit_/μ_nuc_ in 121 multilocus datasets covering 4,676 animal species and found that μ_mit_/μ_nuc_ varies widely across taxonomic groups. In vertebrates, μ_mit_/μ_nuc_ is typically above 10 and averages around 20. Invertebrates tend to have much lower μ_mit_/μ_nuc_ values, ranging from 2 to 6. Across crustaceans, μ_mit_/μ_nuc_ estimates range from 2.0 to 10.4 with an average of 5.9. We estimated μ_mit_/μ_nuc_ using our genome-wide μ estimate and a μ_mi_ estimate of 1.1E-8 substitutions/site/year for the mitochondrial CO1 gene, derived from the most closely related transisthmian species pair (*A. colombiensis/A. estuariensis).* Our estimate of μ_mit_/μ_nuc_ was 4.3, a value very similar to the average value observed across other crustacean groups. Several factors have been used to explain the lower μ_mit_/μ_nuc_ in arthropods including a lower mass-specific metabolic rate (Makarieva et al. 2008), taxonomic differences in the ratio of mtDNA/nuDNA replication cycles per generation (Mishra and Chan 2014), and a negative correlation between per generation mutation rates and effective population size (Lynch 2010).

SNPs identified in our GBS data can provide a rough approximation of DNA substitution types and the ratio of transitions (Ts) to transversions (Tv) in *Alpheus* (Fig. 1). However, unlike experimental assays of mutation rate, our analysis cannot determine the directionality of substitution types. The stringency for missing data and taxa inclusion influences our estimates of Ts/Tv slightly (Supp. Table 1), an effect that has been observed in other reduced representation sequencing datasets (Shafer et al. 2017). Our primary dataset, Am4s2, had a Ts/Tv ratio of 1.399, while the more conservative Am3s3 dataset had a Ts/Tv ratio of 1.489. These Ts/Tv ratios are comparable to the Ts/Tv ratio of 1.22 in *Drosophila* (Petrov and Hartl 1999) but lower than the accepted Ts/Tv ratio of ~2.1 for humans (Ebersberger et al. 2002; DePristo et al. 2011). Variation in Ts/Tv ratios may be due to fundamental differences in point substitution processes (Keller et al. 2007) or an artifact of the sequencing approach (Davey et al. 2013; Ba et al. 2017).

### Locus Bias in GBS Data

Our analyses also examined the effects of molecular divergence on locus recovery from GBS data, providing empirical evidence for a phylogenetic signal in the distribution of missing data when reduced representation methods are applied to species-level investigations. This finding has important implications for establishing unbiased filtering parameters in bioinformatic pipelines, as data from GBS and restriction-site associated DNA (RAD) sequencing are now commonly used for demographic and phylogenomic analyses at both shallow and deep time scales (Andrews et al. 2016). Because these methods utilize restriction enzymes to reduce the genome, they require conservation of enzyme recognition cut sites to recover shared data among individuals. In theory, a mutation in a cut site would result in either allelic dropout at shallow time scales and bias population genetic inferences (Gautier et al. 2013), or potentially the loss of phylogenetically informative loci at deeper timescales. Missing data can also arise from low or uneven sequencing coverage across samples (Eaton et al. 2017). Simulations of GBS datasets at phylogenetic scales found that bioinformatic filtering to reduce missing data can select for loci with lower mutation rates that are more likely to be genotyped across taxa (Huang and Knowles 2016). Determining the cause of missing data in a GBS dataset can help inform bioinformatic processing decisions, which can, in turn, influence downstream phylogenetic and population genetic inferences (Shafer et al. 2017; Díaz-Arce and Rodríguez-Ezpeleta 2019).

Using linear models and ordination we demonstrated that the amount of missing data between samples was strongly influenced by phylogenetic relatedness and, to a lesser extent, sequencing depth. This result suggested that a strict missing data filter may impose a phylogenetically-informed bias on the retained data. It is likely that strict filtering parameters will preferentially retain phylogenetically conserved loci that are subject to purifying selection. We examined the proportion of loci shared across three or more species for putative protein coding and non-coding loci; the proportion of loci recovered from three or more species was more than 30% higher for coding than non-coding tags (7.1% and 5.4%, respectively). We also tested the influence of filtering parameters on demographic parameter estimates obtained from G-PhoCS; a total of 10 different datasets were applied that varied in the amount of missing data allowed across individuals and/or species. Our most conservative dataset, requiring a locus to be present in at least three species, produced significantly lower divergence time estimates for all three species pairs. This result suggests that a stringent missing data filter selects for loci that are more conserved across species and therefore have lower substitution rates, supporting simulation findings (Huang and Knowles 2016). Strict filtering thresholds also impacted other demographic parameters, such as estimates of current and ancestral effective population size (Θ)(Supp. File 2). Other studies that have used GBS or RAD datasets for interspecific demographic modelling often only include loci found in all study species (Campagna et al. 2015; Oswald et al. 2017). Our results highlight the risk of using strong filters for missing data with G-PhoCS and other demographic methods, and instead suggest demographic models using reduced representation methods should be tested with a range of missing data.

### Divergence Time Estimates

The sequence of divergence times for *Alpheus* transisthmian species pairs based on our GBS dataset is largely consistent with earlier, coalescent based estimates based on sequence data from nuclear protein coding genes. Model-based estimates of τ using the software IMa (Hey and Nielsen 2004) and MCMCCoal (Yang 2002) on eight nuclear genes found that *A. panamensis/A. formosus* was the first species pair to be separated, followed by *A. malleator/A. wonkimi,* with the most recent split being between *Alpheus colombiensis/A. estuariensis;* the 95% confidence intervals of τ for these latter two species pairs overlapped considerably (Hurt et al. 2009). In this study of the broader nuclear genome (Table 2), we again found *A. panamensis* and *A. formosus* diverging first at an estimated T = 4.73 Ma, followed by *A. colombiensis* and *A. estuariensis* (T = 3.0 Ma), with *A. wonkimi* and *A. malleator* the most recently diverged pair (T = 2.25 Ma). It is intuitive that these often intertidal species would be clustered and recent in their divergence times as they would have had more opportunities for dispersal during the final stages of the formation of the Isthmus than species inhabiting rocky intertidal habitats (O’Dea et al. 2016).

Mitochondrial loci evolve separately from nuclear genes, providing an independent dataset for examining divergence. K2P pairwise sequence distances in the mitochondrial COI barcoding gene have indicated a partially different order of divergence, showing *A. malleator/A. wonkimi* diverging first (K2P = 11.5%), with *A. panamensis/A. formosus* diverging next (K2P = 9.5%), and *A. colombiensis/A. estuariensis* as the last-diverging pair (K2P =6.8%) (Lessios 2008). Coalescent based divergence times account for polymorphisms within ancestral taxa which can influence split estimates while K2P distances do not, thus the difference in divergence order between these two approaches may reflect ancestral lineage sorting.

### Mutation rates, vicariance patterns, and the timing of final closure of the Isthmus of Panama

Comparison of our estimate of μ to other established mutation rate estimates can provide insight for the ongoing debate surrounding the timing of the closure of the Isthmus of Panama. We compared our estimate for μ when using a calibration point of divergence at the more broadly supported estimate of 3 Ma (O’Dea et al. 2016) vs. the suggestion of a considerably older timing of 10 Ma (Sepulchre et al. 2014, Bacon et al. 2015a; Montes et al. 2015). We found that the latter resulted in an almost 5-fold lower estimate of mutation rate than the rates found for *Drosophila, Daphnia,* and other multicellular eukaryotes (Denver et al. 2009; Keightley et al. 2009; Ossowski et al. 2009; Keith et al. 2016). The estimate of μ_mit_ is also consistent with estimates from independently calibrated arthropod taxa when calibrated with a 3 Ma Isthmus. While it is possible that *Alpheus* have a considerably lower nuclear mutation rate than other studied taxa, it is unlikely that both nuclear and mitochondrial genomes would exhibit unusually low mutation rates as these processes occur independently (Knowlton and Weigt 1998).

Not all transisthmian species pairs reflect recent, clustered vicariant events (e.g., three of eight *Alpheus* pairs studied by Hurt et al. (2009) and some other taxa reviewed in O’Dea et al. (2016)). However, the contemporaneous divergence time estimates of multiple transisthmian species pairs (Hurt et al. 2009; O’Dea et al. 2016) supports the utility of the formation of the Isthmus as a calibration point for evolutionary histories. Overdispersion of pairwise distance estimates in mitochondrial genes has been used to refute the established timeline for completion of the Isthmus (Bacon et al. 2015a). However, findings from our GBS dataset that account for polymorphisms in ancestral populations and post-divergence gene flow suggest that divergence times for multiple species pairs occurred within a narrow window at about 3 Ma. Our results support accounting for accurate taxon sampling and coalescent processes in ancestral populations when examining transisthmian speciation events (Marko et al. 2015).

Once formed, the Isthmus of Panama represented an impenetrable barrier for shallow water marine species to migrate between the eastern Pacific and western Atlantic, but opportunities for migration may have fluctuated as closure neared final completion. Results from our G-PhoCS analyses largely support a complete isolation of eastern Pacific and western Atlantic *Alpheus* populations following the closure of the CAS. Of the six migration parameters estimated, significant post-divergence gene flow was only found for *A. panamensis/A. formosus,* the species pair with the oldest divergence time (approx. 4.73 Ma). Best estimates of migration rates in this pair were exceedingly low; while gene flow was bidirectional, greater migration was inferred from the eastern Pacific species towards the western Atlantic species; this is consistent with models showing that strong currents passed through the straits from the Pacific into the Caribbean leading up to its final closure (O’Dea 2016; Schneider 2006). *Alpheus panamensis/A. formosus* are both common, free-living species that occupy a wide range of habitats, including under rocks and in rock crevices, in dead and living coral rubble, and in sand/mud mixed substrate. Tolerance to a diversity of habitats may have facilitated trans-oceanic passage during the final stages of Isthmus formation when sub-optimal habitats may have been encountered by migrants, and even today these species are occasionally capable of producing fertile hybrid clutches (Knowlton et al. 1993). We found that the inclusion of post-divergence migration parameters was important for obtaining robust estimates of mutation rates. Failure to account for post-split migration results in a negative bias in estimates of divergence times and inflates estimates of genome-wide substitution rates. For example, the estimated divergence time for *A. panamensis/A. formosus* (τ_PF_) was reduced by 19% when migration was not included in the model (Fig. 3).

## Conclusion

Our results add an additional layer of support for a recent closure of the Panamanian Isthmus which has broad-ranging implications across evolutionary biology. The agreement between our estimate of the genomic mutation rate and experimentally derived mutation rates in multiple model organisms makes an older Isthmus highly unlikely. Though widely criticized (Lessios 2015; Marko et al. 2015; O’Dea et al. 2016), the suggestion that the closure of the Isthmus may have occurred as early as 23 million years ago has cast doubt on decades of studies across multiple disciplines that have relied on the widely accepted closure date of 3 million years (Hoorn and Flantua 2015). The multi-locus approach employed here highlights the importance of accounting for ancestral lineage sorting when using geological events to calibrate molecular processes.

The nuclear mutation rate and evidence for phylogenetic signal in loci identified here can inform studies using reduced representation methods to address phylogeographic and demographic histories in non-model taxa. Our empirical-based estimate of the nuclear mutation rate aligns well with experimentally determined mutation rates in model arthropod species suggesting that these rates may be applied more broadly to studies in other taxonomic groups. Results from our exploration of filtering parameters serve as a cautionary tale for the adherence to strict bioinformatic filtering parameters. To our knowledge, this is the first use of transisthmian species pairs to calibrate the rate of molecular evolution from reduced-representation sequencing data.

## Author Contributions

All authors contributed to the design of the study. CH and NK collected tissue samples. JI performed the molecular lab work. KS and CH analyzed the data and drafted the manuscript. All authors provided critical input to the manuscript.

## Acknowledgements

This work was funded by a University of Miami Scientists and Engineers Expanding Diversity and Success (SEEDS) grant (NSF #0820128) and a University of Miami General Research Support Award. KS was funded by the National Science Foundation Graduate Research Fellowship under Grant No. 1545870 and the Department of Education Graduate Assistance in Areas of National Need Fellowship Grant No. P200A150101.

## Data Archiving

Upon acceptance, raw demultiplexed genotype-by-sequencing DNA sequences will be made available on NCBI SRA. CO1 sequences will be made available through Genbank. Input files for G-PhoCS will be available on Dryad. Scripts used for data analysis will be available on the corresponding author’s Github.

