## Supplementary Information for "Base-substitution mutation rate across the nuclear genome of *Alpheus* snapping shrimp and the timing of isolation by the Isthmus of Panama"

**Supp. Table 1.** Description of different datasets used in the paper, including taxa included, the minimum number of individuals at each locus (m), the minimum number of species at each locus (s), the total number of loci, the number of putative neutral loci, the number of loci used in G-PhoCS in cases where loci were randomly subsampled, the average number of individuals per locus in the neutral loci, and the transition/transversion (Ts/Tv) ratio.

| **Dataset** | **Taxa** | **m** | **s** | **Loci : Total / Neutral / Used in G-PhoCS** | **Avg. individuals per locus** | **Ts/Tv of neutral loci** |
|---|---|---|---|---|---|---|
| Am4 | All | 4 | 1 | 56,838 / 52,923 / NA | 5.17 | 1.398 |
| Am4s2 | All | 4 | 2 | 48,062 / 44,960 / 14,986 | 5.34 | 1.399 |
| 23m6s2 | All except one poorly sequenced *A. malleator* | 6 | 2 | 19,276 / 18,230 / 18,230 | 7.23 | 1.426 |
| Am3s3 | All | 3 | 3 | 3,481 / 3,475 / 3,475 | 7.45 | 1.489 |
| PFECm3s2 | Excluding *A. malleator* and *A. wonkimi* | 3 | 2 | 66,235 / 62,791 / 20,930 | 4.51 | 1.378 |
| EC_Fm3s2 (*a*) | *A. estuariensis / A. colombiensis A. formosus* | 3 | 2 | 28,904 / 27,589 / 27,589 | 4.23 | 1.359 |
| EC_Wm3s2 (*b*) | *A. estuariensis / A. colombiensis A. wonkimi* | 3 | 2 | 29,092 / 27675 / 27675 | 4.26 | 1.357 |
| WM_Pm3s2 (*a*) | *A. wonkimi / A. malleator A. panamensis* | 3 | 2 | 9,323 / 8,835 / 8,835 | 4.23 | 1.538 |
| WM_Cm3s2 (*b*) | *A. wonkimi / A. malleator A. colombiensis* | 3 | 2 | 7,301 / 7,058 / 7,058 | 3.56 | 1.594 |
| PF_Em3s2 (*a*) | *A. panamensis / A. formosus A. estuariensis* | 3 | 2 | 37,546 / 34,989 / 34,989 | 4.68 | 1.392 |
| PF_Wm3s2 (*b*) | *A. panamensis / A. formosus A. wonkimi* | 3 | 2 | 38,315 / 36,218 /18,108 | 4.79 | 1.401 |

**Supp. Table 2.** Configuration parameters for G-PhoCS, including priors for estimating effective ancestral population size (Θ), absolute effective ancestral population size in number of individuals ($N_e$), and divergence times (τ) for different nodes of the phylogeny (WM=*A. wonkimi*/*A. malleator*, EC=*A. estuariensis*/*A. colombiensis*, PF=*A. panamensis*/*A. formosus*, PFWM=common ancestor of PF and WM).

| **Parameter** | **Value** |
|---|---|
| Burn-in iterations | 100,000 |
| Sample iterations | 200,000 |
| Sample-skip | 10 |
| Prior for scaled locus-specific mutation rate (when not constant) | Dirichlet (=1) |
| Prior for all Θ parameters | Gamma (=1, =1000) |
| Prior for all migration rate parameters | Gamma (=0.002, =0.00001) |
| Prior for $\tau_{WM}$, $\tau_{PF}$, $\tau_{EC}$ | Gamma(=1, =300) |
| Prior for $\tau_{PFWM}$, $\tau_{root}$ | Gamma(=1, =100) |
| Initial value to start MCMC for $\tau_{WM}$, $\tau_{PF}$, $\tau_{EC}$ | 0.003 |
| Initial value to start MCMC for $\tau_{PFWM}$, $\tau_{root}$ | 0.01 |

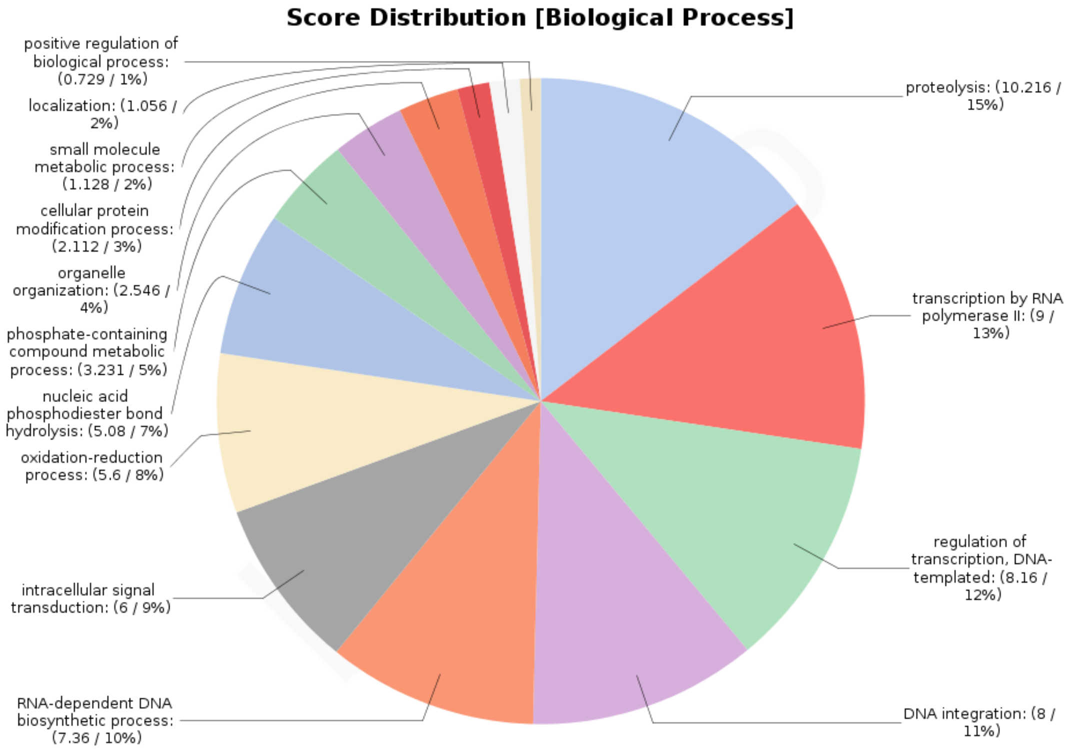
Score Distribution [Biological Process]
positive regulation of biological process: (0.729 / 1%)
localization: (1.056 / 2%)
small molecule metabolic process: (1.128 / 2%)
cellular protein modification process: (2.112 / 3%)
organelle organization: (2.546 / 4%)
phosphate-containing compound metabolic process: (3.231 / 5%)
nucleic acid phosphodiester bond hydrolysis: (5.08 / 7%)
oxidation-reduction process: (5.6 / 8%)
intracellular signal transduction: (6 / 9%)
RNA-dependent DNA biosynthetic process: (7.36 / 10%)
proteolysis: (10.216 / 15%)
transcription by RNA polymerase II: (9 / 13%)
regulation of transcription, DNA-templated: (8.16 / 12%)
DNA integration: (8 / 11%)

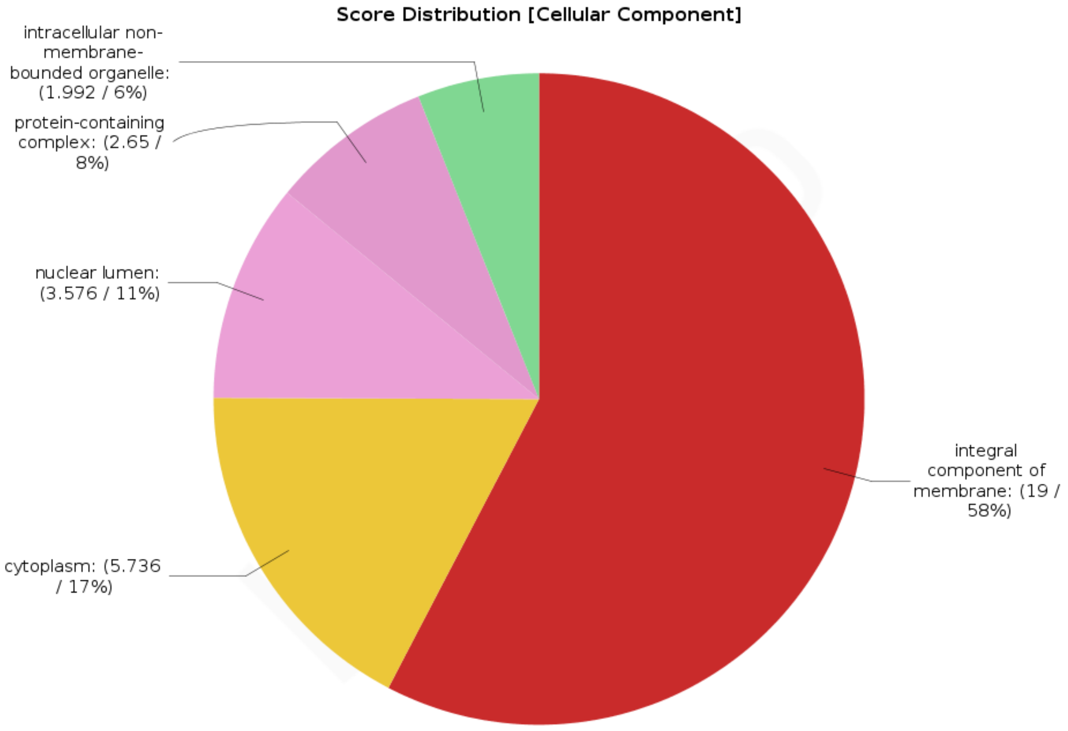
Score Distribution [Cellular Component]
intracellular non-membrane-bounded organelle: (1.992 / 6%)
protein-containing complex: (2.65 / 8%)
nuclear lumen: (3.576 / 11%)
cytoplasm: (5.736 / 17%)
integral component of membrane: (19 / 58%)

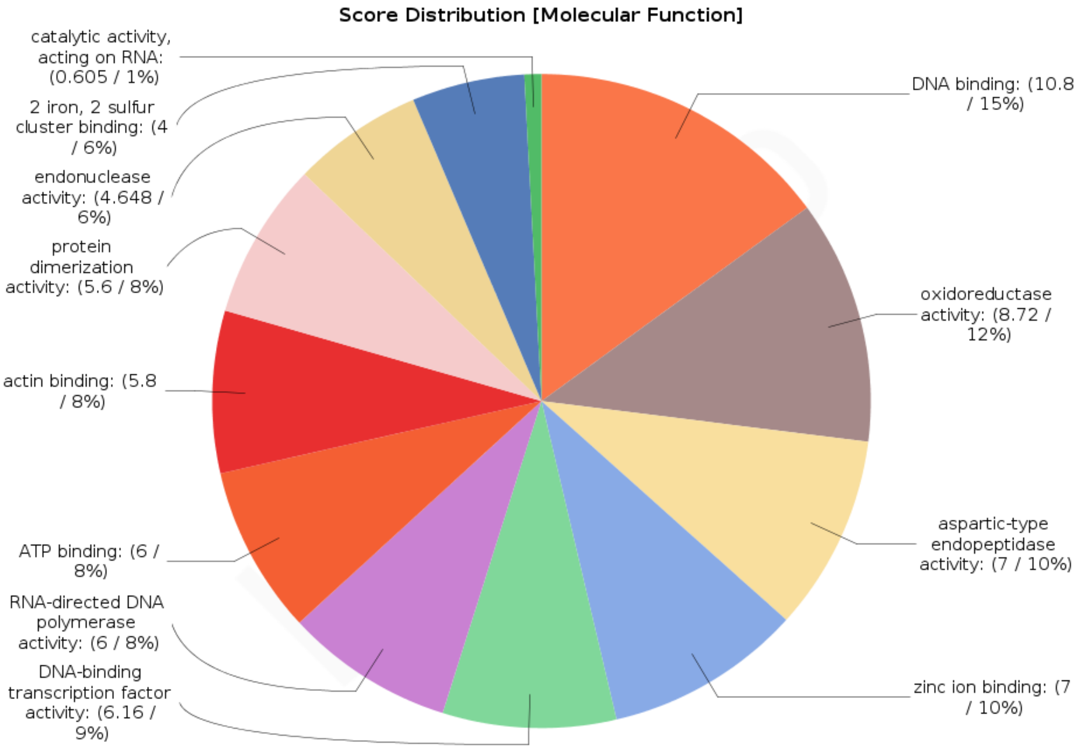


**Supp. Figure 1.** Distribution of gene ontology (GO) annotations from Blast2GO for putative protein-coding loci. In parentheses for each GO term is the Node Score (calculated as the sum of sequences associated with the GO term weighted by how distantly related the GO annotation of the sequence is to the given GO term), and the % of total annotated sequences represented by the GO term. For a GO term to be included it had to have at least 4 sequences associated with it.

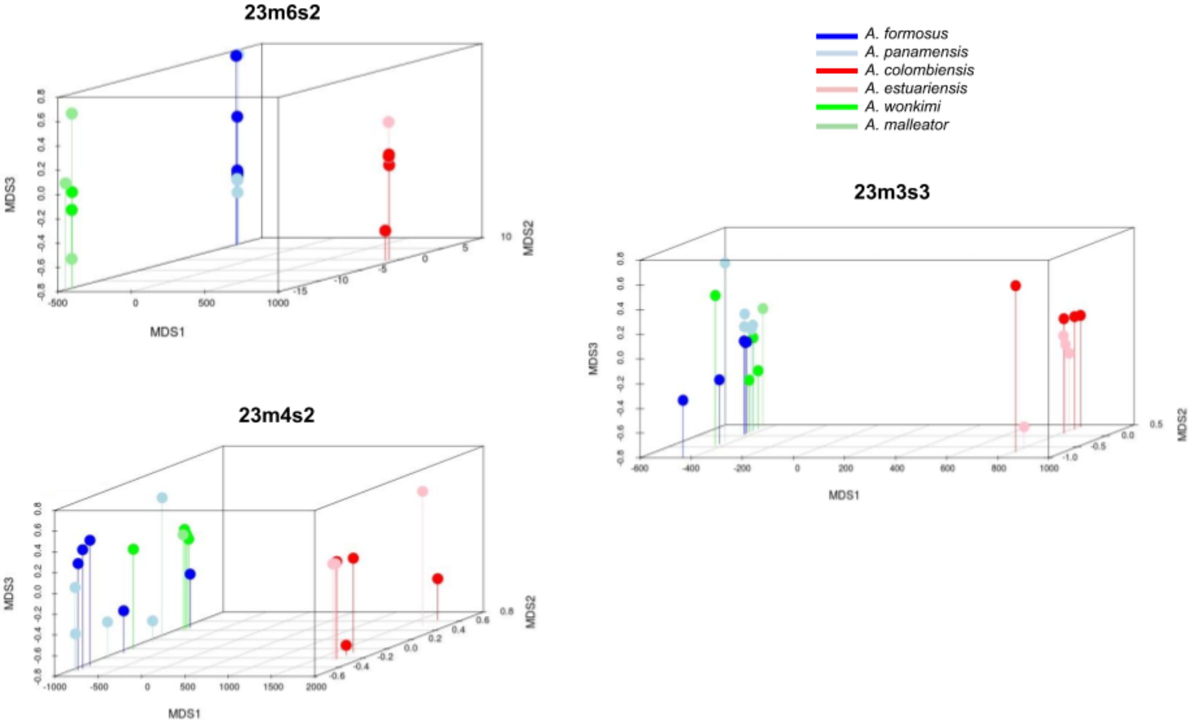


**Supp. Figure 2.** MDS plots with K = 3 ordination of locus-sharing matrix for three datasets varying in minimum number of individuals at each locus (*m*) and minimum number of species at each locus (*s*). Datasets include 23 samples and exclude the one *A. malleator* sample with low sequencing depth. Samples are colored by species.

**A.**

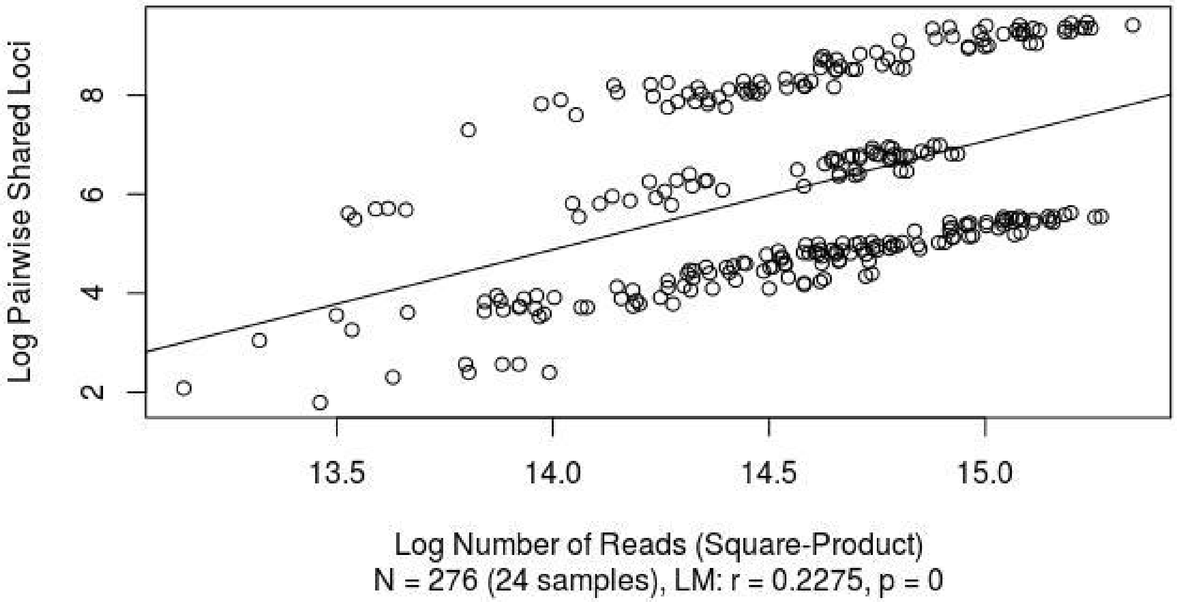


**B.**

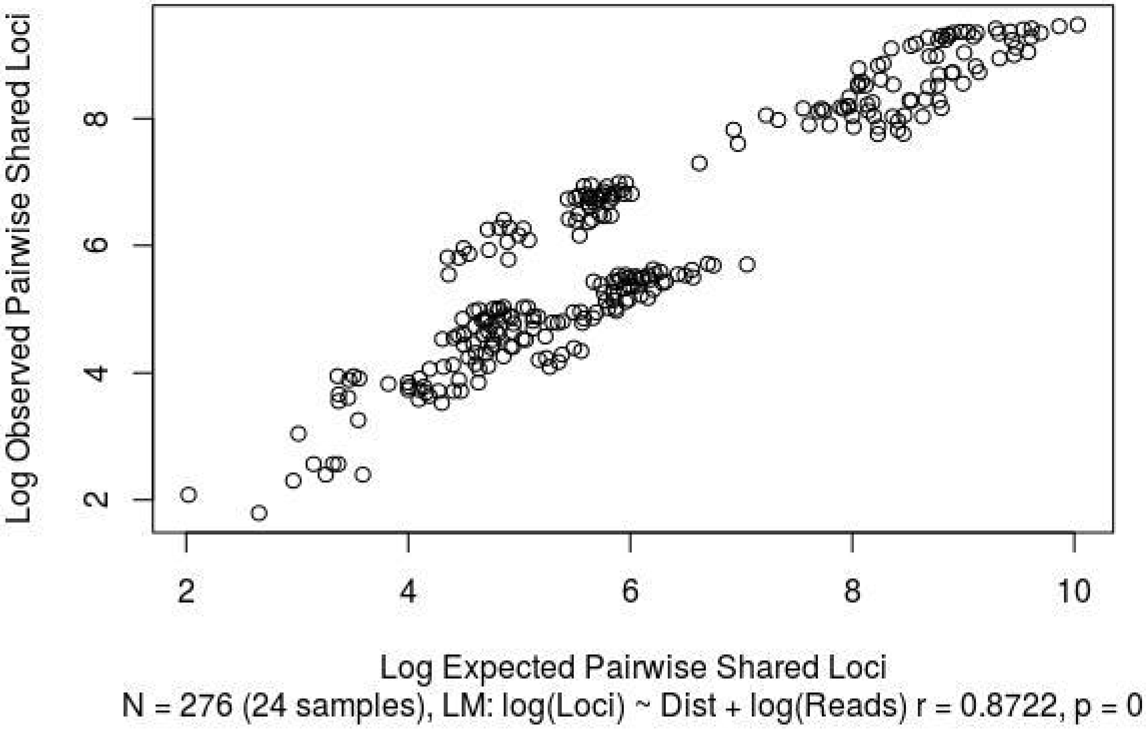


**Supp. Figure 3:** Quantitative modelling of bias in the number of shared loci between samples, using the Am4s2 neutral dataset.. **A.** Log number of pairwise shared loci plotted against the log square-product of the number of sequencing reads for each sample after quality filtering. **B.** Linear model of log expected pairwise shared loci given the phylogenetic distance and log square-product of filtered sequencing reads.
